# A mouse model for cerebral/cortical visual impairment (CVI) impairs vision and disrupts the spatial frequency tuning of neurons in visual cortex

**DOI:** 10.1101/2025.09.19.677390

**Authors:** Dana K. Oakes, Cecilia A. Attaway, Wenxin Zeng, Jun Cai, William Guido, Aaron W. McGee

**Affiliations:** Department of Anatomical Sciences and Neurobiology, School of Medicine; University of Louisville, Louisville, KY, 40202, USA; Department of Ophthalmology and Visual Sciences, School of Medicine; University of Louisville, Louisville, KY, 40202, USA; Pediatric Research Institute, Department of Pediatrics, School of Medicine, University of Louisville, Louisville, KY 40202, USA; Department of Translational Neurosciences, School of Medicine, University of Arizona College of Medicine – Phoenix, Phoenix, AZ, 85004, USA

**Author notes:** Corresponding authors: William Guido, and Aaron W. McGee.

## Abstract

Cerebral/cortical visual impairment (CVI) is a visual disorder associated with perinatal hypoxic injury. The pathophysiology of CVI is poorly understood in part because of the lack of an animal model. Here we developed a murine model of CVI from existing rodent early postnatal hypoxia models for periventricular leukomalacia. Exposure to hypoxia during the equivalent to the human third trimester did not perturb motor function but caused severe impairments in binocular depth perception and visual acuity. Impaired vision was associated with normal retinal function, but reduced size of the visual thalamus, and aberrant tuning for spatial frequency by populations of excitatory neurons in primary visual cortex. This murine model of CVI provides a framework for triangulating circuit deficits with specific visual impairments and testing potential therapeutic interventions.

## Introduction

Cerebral/cortical visual impairment (CVI) is the leading cause of visual impairment in children of developing nations, affecting as many as 1 in 30 ages 5 to 11 (Hatton et al. 2007; Williams et al. 2021). In the U.S., CVI is the most prevalent diagnosis of children with visual impairment younger than 3 years of age (24%) (Hatton et al. 2007). Diagnosis of cerebral/cortical visual impairment (CVI) is increasing, likely due to improved survival of infants with perinatal neurologic injury (Chokron & Dutton 2016; Good et al. 2001; Hoyt 2003; Huo et al. 1999). Despite such prevalence, treatment of CVI is an unmet clinical need.

CVI is caused by damage to the central visual pathways and results in a spectrum of visual deficits. Patients with CVI may exhibit low visual acuity, decreased in visual attention, visual field defects, oculomotor dysfunction, and/or reduced contrast sensitivity, as well as more complex visual impairments (Dutton 2006; Fazzi et al. 2007; Good et al. 2001; Roman et al. 2010). CVI is associated with a variety of perinatal defects including congenital central nervous system malformation, traumatic brain injury, cerebral vascular accident, metabolic disorders, meningitis/encephalitis, and severe epilepsy (Huo et al. 1999). However, CVI is most often associated with perinatal hypoxia (Huo et al. 1999; Jacobson & Dutton 2000; Pagliano et al. 2007), and is common in cases of cerebral palsy and preterm birth (Geldof et al. 2015; Macintyre-Béon et al. 2013).

Perinatal hypoxia also causes periventricular leukomalacia (PVL), injury to the white matter surrounding the ventricles that is characterized by reactive gliosis and focal necrosis (Folkerth 2006). PVL is also closely associated with preterm birth and CVI (Folkerth 2006; Geldof et al. 2015). Between 66 to 94 percent of patients with PVL are also diagnosed with CVI (Bauer & Papadelis 2019; Cioni et al. 1997). Periventricular white matter contains projections from thalamus to cortex, and loss of these projections are hypothesized to be the principal cause of the visual deficits in CVI. Nonetheless, the perturbations to visual circuitry underlying visual deficits in CVI are poorly understood.

A model of CVI is necessary to improve understanding of this disorder and guide development of effective treatment. To date, no animal model has been developed for CVI although numerous features of the maturation and function of visual circuitry are conserved between primates, cats, and rodents (Huberman & Niell 2011; Priebe & McGee 2014). However, there are rodent models of PVL (Turner et al. 2003; Weiss et al. 2004). These models take advantage of the altricial nature of rodent postnatal development where the first two weeks postnatal are equivalent to human third trimester development (Almeida et al. 2020; Ment et al. 1998). These models of PVL mimic perinatal hypoxia by exposing dams with pups to durations of lower oxygen levels (9.5%) before returning them to normoxic (18%) conditions. Mice or rats are exposed to this early postnatal hypoxia for 7 to 30 days beginning at postnatal (P) day 3 (Ment et al. 1998; Turner et al. 2003; Weiss et al. 2004). Here we adapted the early postnatal hypoxia mouse model of PVL into an effective model for CVI.

## Results

To assess the effectiveness of exposure to early postnatal hypoxia as a model for CVI, first we performed a battery of behavioral tests to measure motor and visual function on adult mice that were exposed to hypoxic conditions beginning at P3 for 7 days (d), 14d, or 30d, and then returned to normal housing (Fig. 1). Weeks later, these cohorts were then compared to normoxic controls. Behavioral tests included the rotarod task to evaluate motor performance and learning, the pole descent cliff task (PDCT) to measure binocular depth perception, and the visual water task to measure acuity (Boone et al. 2021; Fox 1965; Jones & Roberts 1968; Park et al. 2014; Prusky et al. 2000; Stephany et al. 2014) (Fig. 2). Then we examined the physiologic function of the retina with electroretinograms (Fig. 3), the anatomical organization of visual circuitry with anterograde labelling of retinofugal projections (Fig. 4 and 5), and the function of populations of neurons in primary visual cortex with *in vivo* calcium imaging at neuronal resolution (Figs. 6-8).

**Figure 1.**
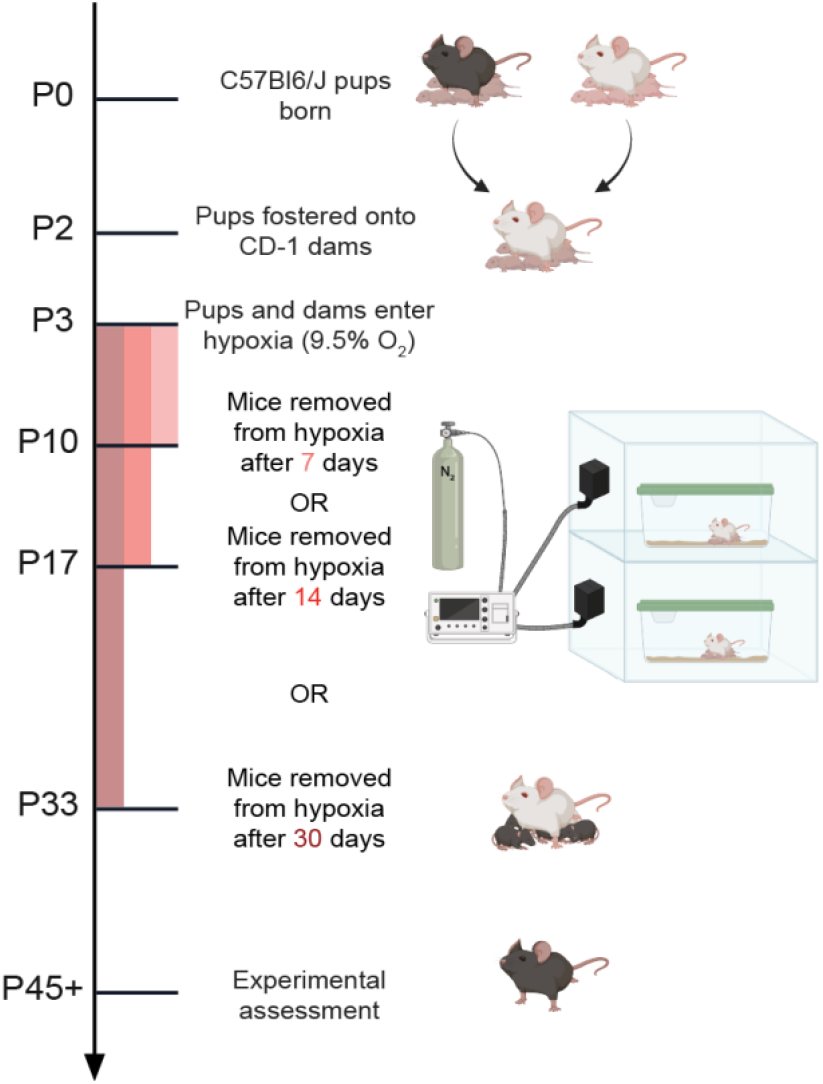
Experimental approach and timeline. A. C57BL/6J pups are fostered onto CD1 dams at postnatal day (P) 2. At P3, the litters and dams are placed in hypoxia chambers (9.5 ± 1% oxygen) until P10 (7 days, pink), P17 (14 days, red), or P33 (30 days, dark red) and then returned to normoxic conditions. Experimental assessments were performed on mice aged at P45 or older.

**Figure 2.**
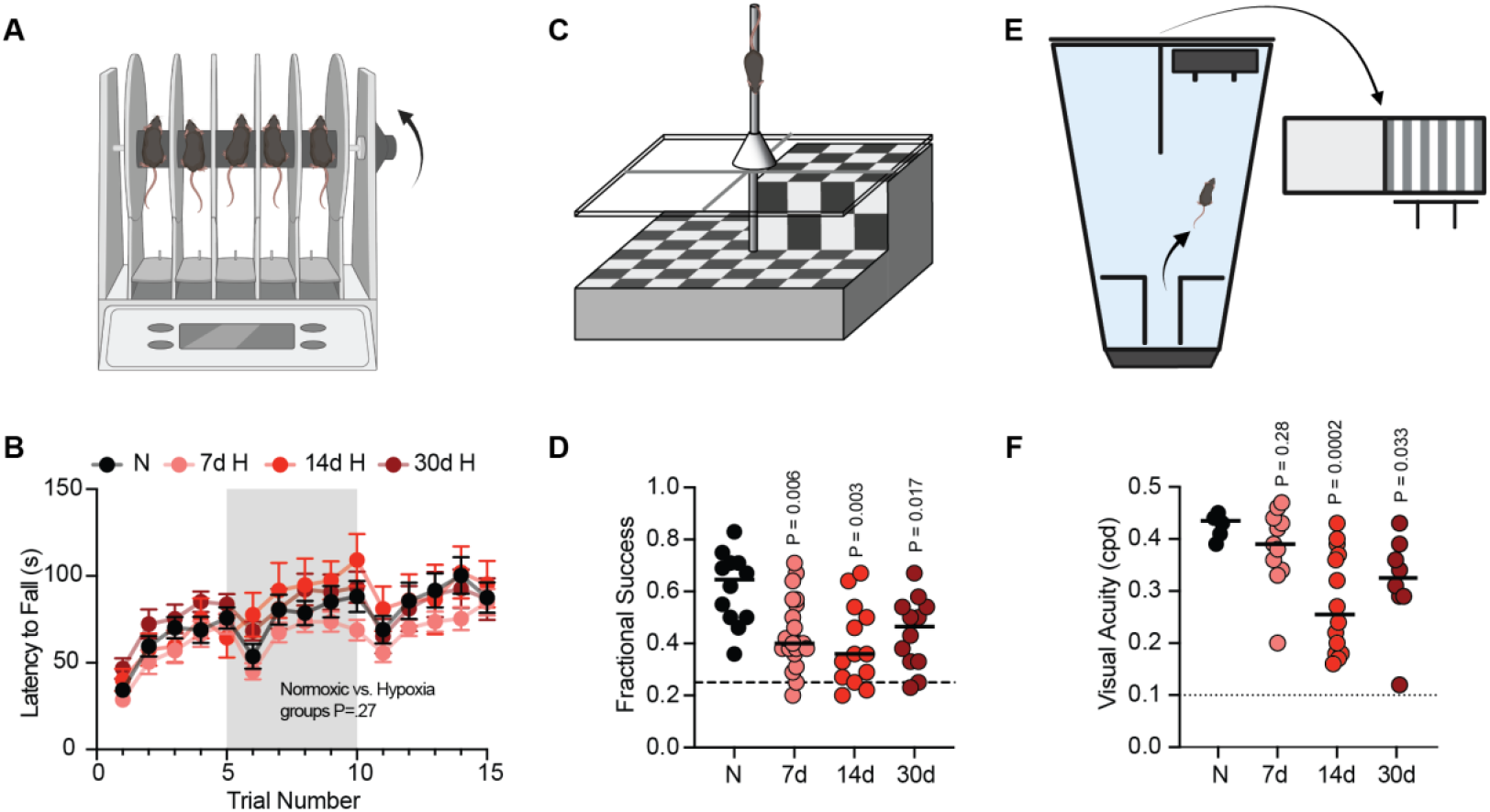
Motor performance and visual function of normoxic controls and mice receiving early postnatal hypoxia. **(A)** Schematic of the rotarod apparatus. **(B)** Latency to fall for normoxic mice (N, n=17 black) and mice exposed to hypoxic conditions for 7d (7d H, n=18, pink), 14d (14d H, n=9 red) or 30d (30d H, n=11, maroon) days. Mice were tested for 5 trials a day on 3 consecutive days. Each point represents the group mean +SEM. Grey and white sections delineate days of testing. F (3, 51) = 1.363, P=.27, Mixed effects analysis for normoxic vs hypoxia exposure. **(C)** Schematic of pole descent visual cliff task **(D)** Fractional success rates of normoxic (N) (n=12, black circles), 7d (n=21, pink), 14d (n=13, red), and 30d (n=12, maroon) hypoxic (H) mice. Circles represent individual mice. Horizontal lines are means. The dashed line indicates chance success (0.25). Comparisons between normoxic and each hypoxic group (7d, P=.006; 14d, P=.003; 30d, P=.017, Welch’s ANOVA) **(E)** Schematic of the visual water task (sinusoidal not square gratings as depicted) **(F)** Visual acuity in cycles per degree (cpd) for normoxic (N, n=6), 7d hypoxia (7d, n=11), 14d hypoxia (14d, n=14), and 30d hypoxia mice (30d, n=8). Circles represent individual mice. Dotted line represents the minimum training threshold (0.1 cpd) for testing. Horizontal lines are means. Comparisons between normoxic and hypoxic groups (7d, P=.28; 14d, P=.0002; 30d, P=033, Welch’s ANOVA).

**Figure 3.**
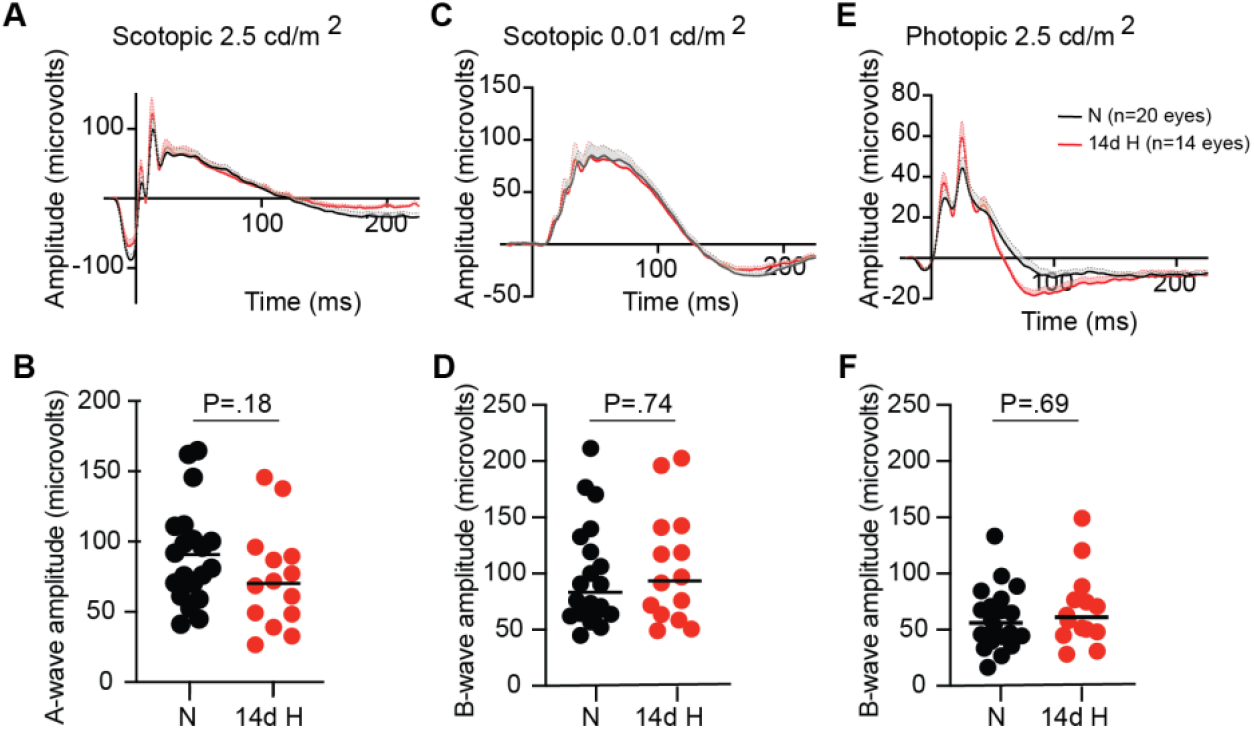
Full field ERGs are normal in 14d hypoxia mice. Average ERG waveforms for adult normoxic control mice (N, n=20 eyes) and adult 14d hypoxia mice (14d H, n= 14 eyes). **(A**,**B)** Scotopic flash waveform and measurement of the a-wave amplitude. P =.18, Welch’s (W) t-test. **(C**,**D)** Scotopic flash waveform and measurement of the b-wave amplitude. P =.74, W t-test. **(E**,**F)** Photopic flash waveform and measurement of the b-wave amplitude. P =.69, W t-test.

**Figure 4.**
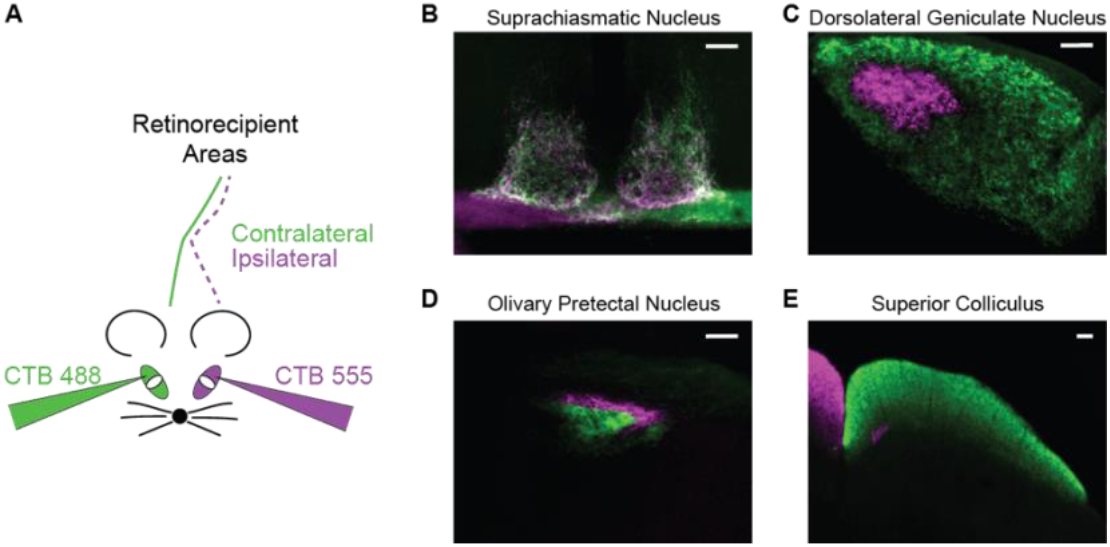
Eye-specific retinofugal projections in normoxic and 14d hypoxic mice. **A**Schematic showing intravitreal injections of cholera toxin b (CTB) subunit conjugated to AlexaFluor 488 or 555. CTB 488 (green) was injected into one eye (contralateral) and CTB 555 (purple) into the other (ipsilateral) eye. **B-E (C)**. Coronal sections through the suprachiasmatic nucleus (B), dorsolateral geniculate nucleus of the thalamus, olivary pretectal nucleus (D) and the superior colliculus (E). CTB terminal fields labeled in green originate from the contralateral eye while those in purple arise from the ipsilateral eye. Scale bars = 100 μm.

**Figure 5.**
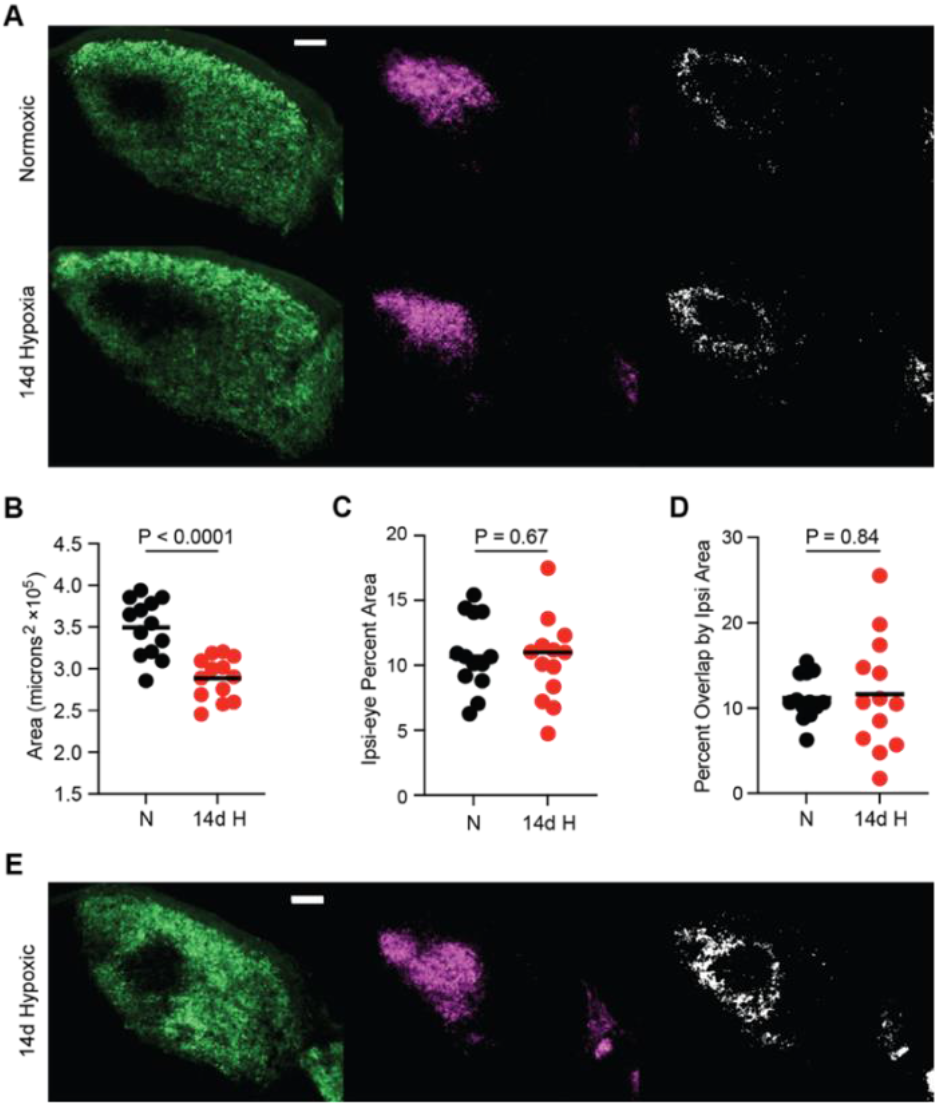
Eye specific patterns of retinal inputs in the dLGN of normoxic and hypoxic mice. **(A)** Examples of coronal sections through the middle part of dLGN of a normoxic mouse and 14d hypoxia mouse. Shown are the retinal terminals fields for the contralateral (green, left) and ipsilateral (purple, middle) eye. The binarized images (white, right) depict the degree of overlap between the terminal fields for the two eyes. Scale bar = 100 μm. **(B)** Size of the dLGN for normoxic (N) and 14d hypoxia (14d H) mice (in square microns x 10^5^). Each circle represents a hemisphere and the horizonal lines represent the mean (N, n = 13 hemispheres, 10 mice; 14d H, n = 13 hemispheres, 8 mice; P < 0.0001, W t-test). (**C)** The percent area of dLGN innervated by retinal projections from the ipsilateral eye for normoxic (N, black circles) and 14d hypoxia (14d H, red circles) mice plotted in panel B. P= 0.67, W t-test. **(D)** Percent area of the overlapping projections by the area of ipsilateral innervation for normoxic (N, black circles) and 14d hypoxia (14d H, red circles) mice plotted in panel B. P =.84, W t-test. **(E)** An example section from 14d hypoxia mouse with a high degree of overlap for comparison. Contralateral (green, left) and ipsilateral (purple, middle) retinal projections to dLGN (white, overlap). Scale bar = 100 μm.

**Figure 6.**
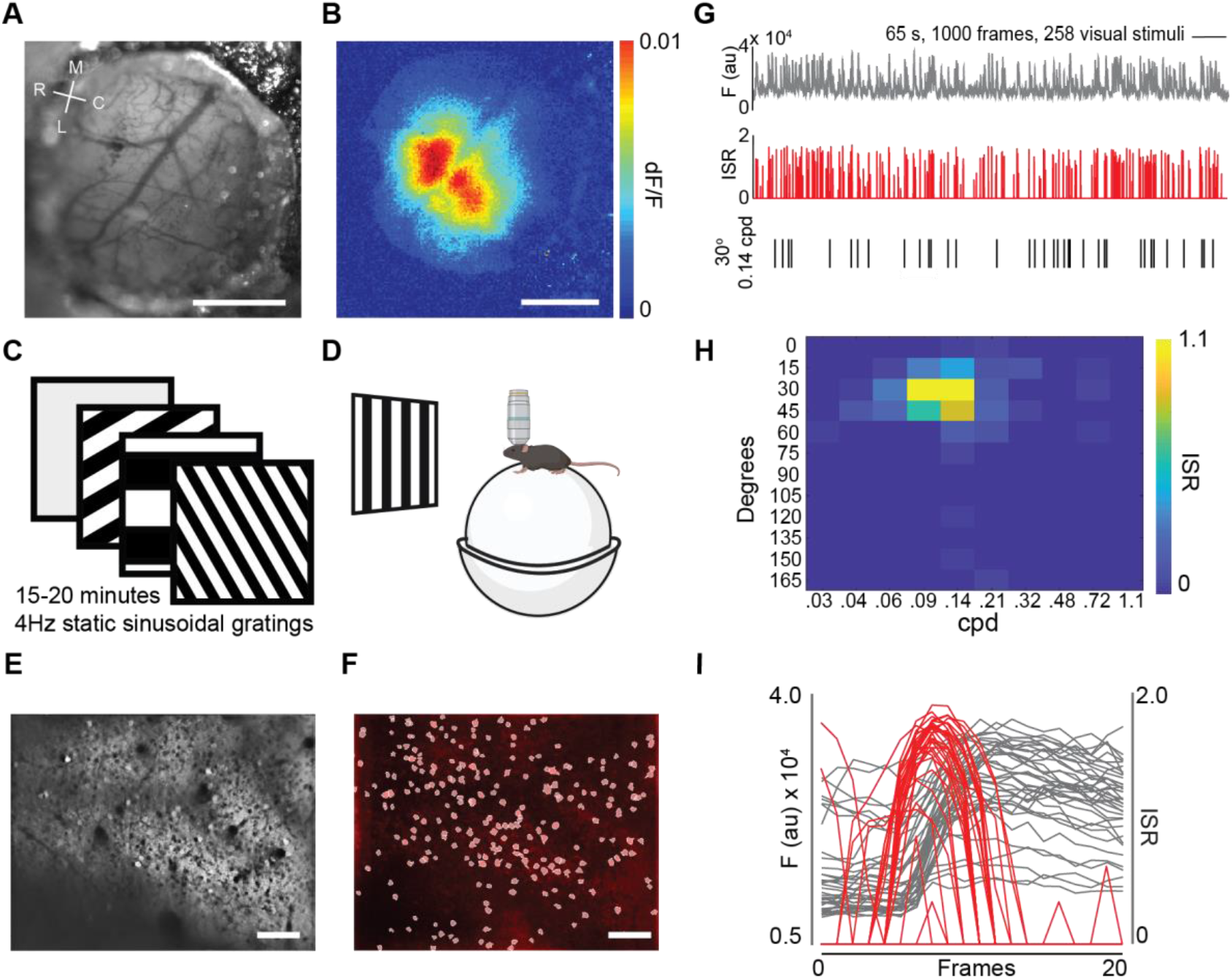
Calcium imaging of primary visual cortex (V1) in an awake head fixed mouse. **(A)** An example of a 3 mm diameter cranial window centered over the visual cortex. Coordinate axes rostral (R) - caudal (C) and medial (M)–lateral (L). Scale bar = 1mm. **(B)** Wide field calcium imaging of neural activity in response to a drifting horizontal bar (20 degrees wide and 2 degrees high). Scale bar = 1 mm. (**C)** Schematic of the visual stimuli. Gratings (sinusoidal not square as depicted) are presented at 15 degrees intervals in orientation between 0.028 and 1.08 cpd spaced at half octaves (log(1.5), along with an isoluminant grey screen in random order at 4 Hz for 15 to 20 minutes. (**D)** Illustration of the calcium imaging set up for alert mice. Mice are head-fixed and freely running on a spherical treadmill floating on a column of air while positioned with a monitor 35cm away centered on azimuth at zero elevation. An occluder (not shown) is placed in front of contralateral or ipsilateral eye in separate imaging sessions. (**E)** An example of the imaging plane in the binocular zone of visual cortex. **(F)** White circles correspond to regions of interest (ROIs) segmented manually from the imaging field in E. Imaging field is 750 μm × 500 μm. Scale bar = 100 μm. **(G)** A trace of fluorescence over time for a region of interest (ROI) / example neuron (grey, top) and the corresponding inferred spike rate (ISR) (red, middle). The timing of presentations of the preferred stimulus (30 degrees, 0.14 cycles per degree (cpd)) during the experiment (black vertical lines, bottom). Images are collected at 15.5 Hz. The scale bar represents 65 seconds, 1000 frames, and 258 visual stimuli (grey horizonal bar, top right). The entire trace represents 15 minutes during which 3,600 gratings were presented from 130 combinations (12 orientations and a grey screen for each of 10 spatial frequencies (SFs). (**H)** Heat map of the ISR for all combinations of orientation and SF (in cpd) from the example neuron in panel G. (**I)** Fluorescent traces (grey lines) superimposed for the 20 frames (1.25 seconds) following the onset of the presentations of the preferred visual stimulus for this example neuron. Red lines represent the positions of inferred spikes.

**Figure 7.**
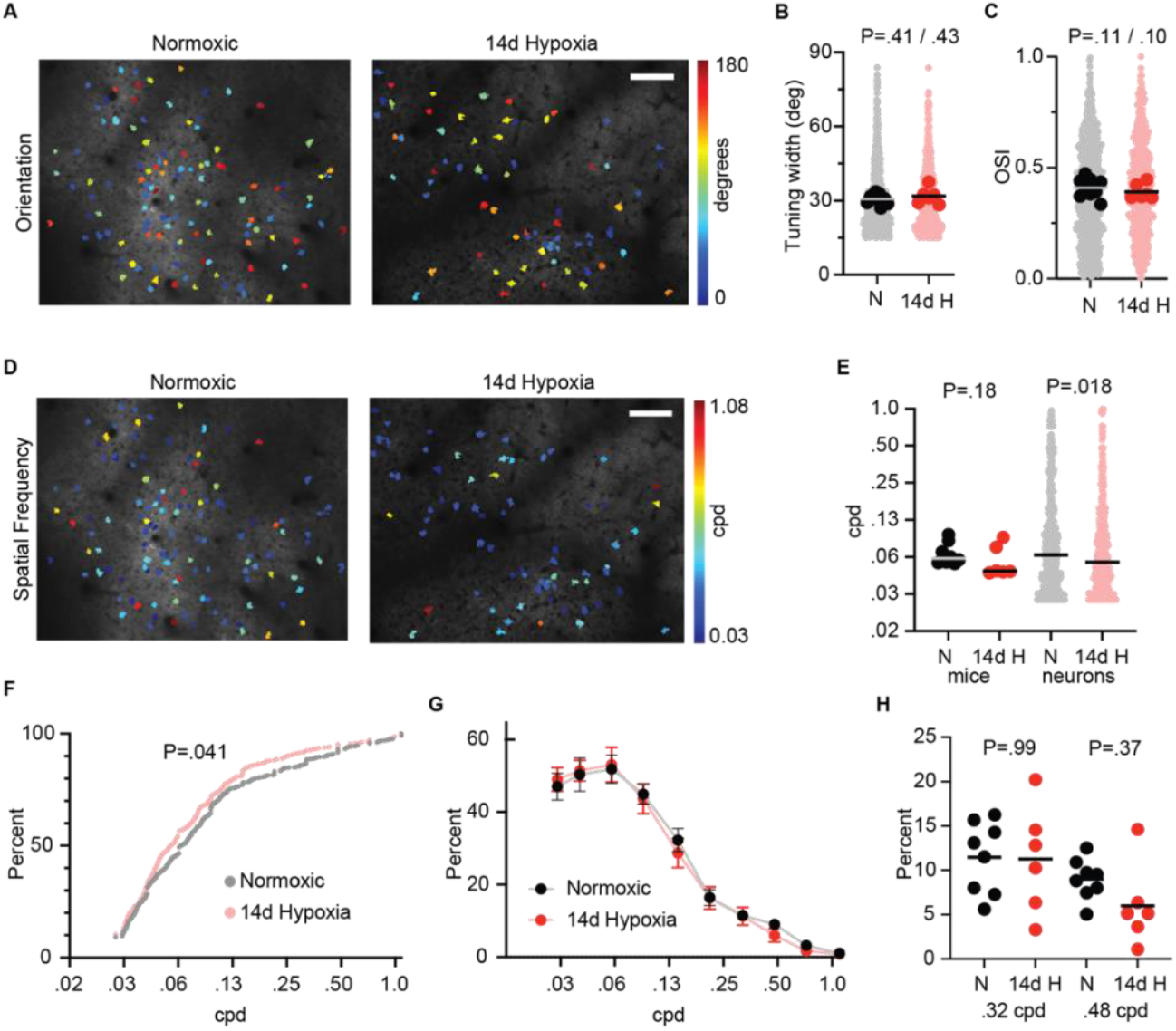
Orientation tuning and spatial frequency tuning of V1 neurons of normoxic and 14d hypoxia mice. **(A)** Example distribution of preferred orientation in image fields of V1 neurons driven by the contralateral eye for a control mouse (Normoxic, left) and a 14d hypoxia mouse (14d Hypoxia, right). Neurons are colored according to their preferred orientation in degrees. Scale bar = 100 μm. **(B)** Half-maximum full tuning width in degrees for normoxic and 14d hypoxia mice (black, n=8; red, n=6) and neurons (grey, n=760; pink, n=516). Comparison of mice, P=.41, and neurons, P=.43, Mann-Whitney (MW) tests (C) Orientation Selectivity Index (OSI) for the same mice and neurons. Comparison of mice, P=.11, and neurons, P=.10, MW tests **(D)** Example distribution of preferred spatial frequency (SF) in cycles per degree (cpd) for the same imaging fields as panel A. **(E)** Preferred SF in cycles per degree (cpd) for the normoxic and 14d hypoxia mice and neurons presented in panel B. Comparison of mice, P=.18, and neurons, P=.018, MW tests. **(F)** Cumulative distribution of the preferred SF for these neurons from normoxic and 14d hypoxia mice (n=760, 516). P=.041, Kolmogorov-Smirnov test. **(G)** Average percent of responsive neurons at each SF presented for normoxic and 14d hypoxia mice (n=8,6). **(H)** Column plot of the percent of responsive neurons for mice in panel G. 0.32 cpd, P=.99; 0.48 cpd, P=.37, Kruskal-Wallis test.

**Figure 8.**
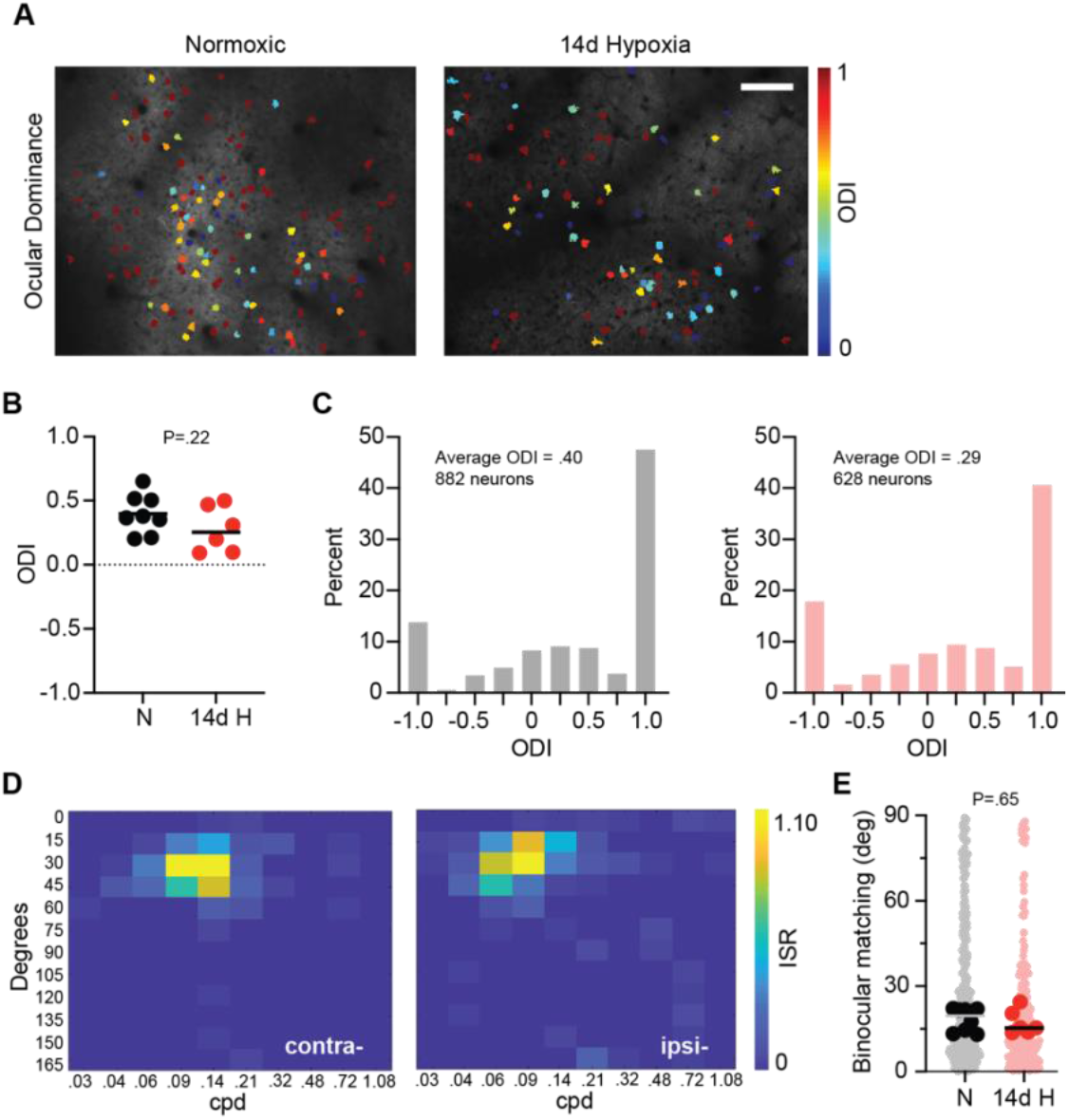
Ocular dominance and binocular matching of orientation preference of V1 neurons for normoxic and 14d hypoxia mice. **(A)** Example distribution of Ocular Dominance Index (ODI) values in image fields of V1 neurons from Fig. 7 panel A, for a control mouse (Normoxic, left) and a 14d hypoxia mouse (14d Hypoxia, right). Neurons are colored according to their ODI value. Scale bar = 100 μm. **(B)** Average ODI values for normoxic (N) and 14d hypoxia (14d H) mice (n=8,6; P=.22, W t-test). **(C)** Histograms of ODI values for neurons from normoxic mice (left, grey bars, n=882) and 14d hypoxia mice (right, pink, n=628). P =.98 between groups, Chi-squared test. **(D)** Heat map of the ISR for all combinations of orientation and SF (in cycles per degree (cpd)) from an example neuron for the contralateral eye (contra-, left) and ipsilateral eye (ipsi-, right) responses. **(E)** Average difference in preferred orientation of binocular neurons for normoxic (N) and 14d hypoxia mice (14d H) (N, black, n=8; 14d H hypoxia, red, n=6, P=.65, W t-test) (N neurons, grey, n=343; 14d H neurons, pink, n=207).

To evaluate if exposure to early postnatal hypoxia perturbs motor function, we tested mice with the rotarod motor performance task (Fig. 2A). The rotarod is a classic measure of gross motor function and motor learning (Jones & Roberts 1968). In this task, mice must balance on a rotating rod while the rod accelerates over time. We measured the latency to task failure for groups of normoxic controls and mice receiving increasing durations of early postnatal hypoxia. Each mouse was tested for 5 trials each day for 3 consecutive days. There was no significant difference in the latency to fall off the rotarod for control mice (n=17) and mice exposed to early postnatal hypoxic conditions for 7d (7d hypoxia) (n=18), 14d (14d hypoxia), (n=9), or 30d (30d hypoxia) (n=11) (Fig. 2B). Mice did not exhibit seizure-like activity or other gross pathophysiological symptoms. This motor performance is consistent with the capacity for mice receiving early postnatal hypoxia to perform visual behavioral assays in the weeks following hypoxia exposure.

Impaired stereopsis is often a symptom of CVI (Dutton et al. 1996). To assess whether early postnatal hypoxia causes deficits in binocular depth perception, we tested mice on the PDCT (Boone et al. 2021) (Fig. 2C). This task is modified from the classic visual cliff task to engage the upper visual field where the binocular overlap is greater in mouse (Boone et al. 2021; Dräger 1975; Fox 1965; Samonds et al. 2019). In each trial, mice descend a pole to towards a glass platform with one of four quadrants positioned closer than the other three quadrants. The success rate represents the number the mouse descends the pole to the quadrant above the closest platform. Normoxic control mice (n=12) displayed a fractional success rate above 60% whereas performance at chance levels is 25% (Fig. 2D). Mice exposed to 7d (n=21), 14d (n=13), or 30d (n=12) hypoxia had significant deficits in this task of depth perception with performance closer to 40% fractional success.

Deficits in visual acuity are a cardinal symptom of CVI (Good et al. 2001; Sakki et al. 2021; Van Genderen et al. 2012). To determine if early postnatal hypoxia is an effective of model of CVI, we measured visual acuity for mice receiving 7d, 14d, or 30d hypoxia with the visual water task (Prusky et al. 2000). The visual water task is an alternative forced choice discrimination swim task in which two visual stimuli are presented at the far end of a trapezoidal tank, one a vertical sinusoidal grating of a user-defined spatial frequency (SF), the other an iso-luminant grey screen. Visual acuity is estimated from the success rate for targeting the side of the tank harboring the hidden platform in front of the SF grating (Fig. 2E). We tested acuity under monocular viewing conditions for the right eye to eliminate potential compensation from binocular vision or differences in acuity between the two eyes resulting from potentially differing perturbations of visual circuitry for each eye.

Normoxic control mice (n=6) possessed typical visual acuity in the range of 0.4 to 0.5 cycles per degree (cpd) (Fig. 2F). By comparison, 7d hypoxia mice (n=11) displayed acuity similar to controls, while mice receiving 14d hypoxia or 30d hypoxia (n=14,8) exhibited significant deficits in acuity. The mean visual acuity of the group of mice receiving 14d hypoxia was near 0.25 cpd, approximately half that of controls, but spanned a range from 0.16 cpd to normal acuity at 0.43 cpd. These deficits in acuity are comparable to those observed in murine models of amblyopia although the range of performance is broader (Prusky & Douglas 2003; Stephany et al. 2018).

Rodent models of PVL expose mice to 7d or 30d hypoxia (Turner et al. 2003; Weiss et al. 2004). We identified similar deficits in binocular depth perception and visual acuity in mice receiving 14d or 30d hypoxia (Fig. 2D,F). Therefore, we chose the 14d hypoxia as a potential model for CVI and investigation of potential visual circuit deficits.

CVI is defined as visual impairment caused by damage to the post-chiasmatic pathways with normal/near normal eye health (Sakki et al. 2018). To determine if retinal function affected by 14d hypoxia, we recorded full-field electroretinograms (ERGs), a direct correlate of the clinical retinal assessment, at flash intensities spanning scotopic to photopic vision (Fig. 3). We examined the wave form and amplitudes of the a-wave and the b-wave for normoxic control mice and 14d hypoxia mice. The overall waveforms of the ERG (Fig. 3 A,C,E), as well as the amplitudes of the a- and b-waves (Fig. 3 B,D,F), were nearly identical between the two groups. Thus, 14d hypoxia does not lead to evident ocular impairment, consistent with the observed deficits in functional vision resulting from disruption of central visual circuitry.

Next, we employed anterograde tracing to visualize the innervation of retinorecipient areas. We performed intravitreal eye injections of cholera toxin subunit B (CTB) conjugated to AlexaFluor dyes into adult mice (Demas et al. 2006; Dilger et al. 2015; Jaubert-Miazza et al. 2005) (Fig. 4A). Retinal projections in 14d hypoxia mice were grossly normal, including to the suprachiasmatic nuclei (SCN) (Fig. 4B), dorsolateral geniculate nucleus (dLGN) (Fig. 4C), olivary pretectal nucleus (OPN) (Fig. 4D), and superior colliculus (SC) (Fig. 4E). Typical overlap of retinal inputs from the two eyes were observed in the SCN while projections to the dLGN, OPN, and SC, displayed little overlap and were dominated by the contralateral eye as expected (Seabrook et al. 2017).

The dLGN is the exclusive relay of visual information to primary visual cortex (V1). Retinal projections from the two eyes overlap at birth but segregate into eye specific domains during early postnatal development (Jaubert-Miazza et al. 2005). This activity-dependent refinement is a hallmark feature of visual system development (Seabrook et al. 2017). Given that this refinement occurs during the period of exposure to hypoxic conditions, we quantified the eye-specific segregation in adult normoxic and 14d hypoxia mice (Fig. 5). Projections from the two eyes remained largely segregated in 14d hypoxia mice, and ipsilateral projections were confined to a patch with a few percent overlap with projections from the contralateral eye along the perimeter of the ipsilateral domain (Fig. 5A). To determine whether hypoxic mice exhibit a greater degree of overlap, we calculated the spatial extent of the dLGN (Fig. 5B), the spatial extent of the ipsilateral eye domain (Fig. 5C), as well as the percent overlap of retinogeniculate projections from the two eyes (Fig. 5D).

The dLGN was nearly 20% smaller on average in 14d hypoxia mice, but the area occupied by the ipsilateral eye domain, and the degree of overlap between the projections from each eye, were similar to normoxic mice (Fig. 5 B-D). However, 14d hypoxia mice exhibited a wider range of variability in the degree of overlap with values spanning 2-25% (Fig. 5 D,E). These higher percentages of overlap were not observed in normoxic mice (Fig. 5 D).

To determine if cortical visual processing is disrupted in 14d hypoxia mice, we performed *in vivo* calcium imaging at neuronal resolution in V1 and calculated the tuning properties for excitatory neurons in Layer (L) 2/3 (Fig. 6). Adult (P45+) normoxic controls and 14d hypoxia mice that expressed the calcium indicator GCaMP6s in forebrain excitatory neurons were implanted with cranial windows over V1 for one hemisphere (Fig. 6A) (Chen et al. 2013; Frantz et al. 2016; Wekselblatt et al. 2016). Next, we mapped the location of the binocular zone of V1 with wide-field calcium imaging with a visual stimulus presenting a drifting white horizonal bar 20 degrees wide and 2 degrees high on a black background (Fig. 6B). Then, we performed calcium imaging at neuronal resolution in the binocular zone while presenting a battery of static sinusoidal gratings that varied in orientation and spatial frequency (SF) (Fig. 6 C,D). Gratings were presented at 4 Hz at random and comprised 12 orientations spaced by 15 degrees across 10 SFs spaced at half octaves from 0.03 cpd to 1.08 cpd, as well as an iso-luminant grey screen. Mice were alert, head-fixed, and positioned on a spherical treadmill during imaging. The visual stimuli were presented to each eye separately by covering the fellow eye with a small occluder.

Regions of interest (ROIs) were segmented by manually by identifying regions with pixel-wise correlation of fluorescence changes across the imaging session (Fig. 6 E,F). The fluorescence signal for each ROI was extracted from this segmentation map by calculating the mean calcium fluorescence within each ROI and then subtracting the median fluorescence from the bordering neuropil (Ringach et al. 2016; Tan et al. 2020) (Fig. 6G). An inferred spike rate (ISR) was derived from the fluorescence signal and employed to determine whether the neuron was visually-responsive (Berens et al. 2018; Brown & McGee 2023; Tan et al. 2020). Tuning properties were calculated from the ISR (Fig. 6 H,I). We compared orientation tuning and SF tuning for neurons responsive to stimulation of the contralateral eye because these groups provided the greatest sampling (Brown et al. 2024; Brown & McGee 2025).

Orientation tuning refines during development following eye opening (∼P14) to reach adult levels near P21 (Hoy & Niell 2015; Tan et al. 2022). There were no evident disruptions of orientation tuning in mice receiving 14d hypoxia. Excitatory L2/3 neurons in V1 displayed a similar distribution of preferred orientation, orientation tuning half-maximum full width, and orientation selectivity index (OSI) (Fig. 7A-C and data not shown). Normoxic control mice and 14d hypoxia mice were similar whether evaluated per animal or by comparing the distribution of neuronal populations (Fig. 7 B,C).

By comparison, SF tuning was somewhat aberrant in 14d hypoxia mice. The median preferred SF for 14d hypoxia mice was lower than the range for normoxic controls for 4 of the 6 14d hypoxia mice, while 2 of the treated mice were within this range (Fig. 7E). These results mirrored the deficits in acuity where most 14d hypoxia mice exhibited lower acuity but a few had normal acuity (Fig. 2E). The distribution of SF tuning by the population of neurons was slightly lower than controls. This deficit was statistically significant when evaluated with a rank-order test (Mann-Whitney) or a test of cumulative distribution (Kolmogorov-Smirnov) (Fig. 7 E,F). To evaluate the distribution of SF tuning per mouse, we plotted for each animal the percentage of neurons that were visually-responsive at each SF tested with a focus on the higher SFs near the normal acuity threshold (Fig. 7G,H). The percent of responsive neurons was similar between groups for 0.32 cpd but the distribution at.48 cpd mirrored the lower median SF values for CVI mice with 5 of the 6 mice exhibiting a percent of responsive neurons at this higher SF that were more than a standard deviation below the mean of the normoxic group (9%±2%).

Last, we assessed whether tuning for binocularity was abnormal in mice receiving 14d hypoxia. The Ocular Dominance Index (ODI) is a measure of the binocularity for neurons (Cang et al. 2005). In mouse V1, a majority of excitatory neurons are predominantly monocular and respond to visual stimuli presented to the contralateral eye (ODI = 1), fewer neurons are binocular (ODI between 1 and -1), and some neurons are predominantly monocular for the ipsilateral eye (ODI = -1) (Brown & McGee 2023; Salinas et al. 2017; Tan et al. 2020). Unlike primates and cats, rodents do not possess ocular dominance columns (Dräger 1975; Grinvald et al. 1986; Hubel et al. 1977; Hubel & Wiesel 1962). Neurons are not spatially organized by binocularity in mouse V1, although a recent study has reported some clustering of neurons preferring the ipsilateral eye (Goltstein et al. 2025) (Fig. 8A). Normoxic control mice and 14d hypoxia mice displayed average ODI values per mouse that were similar (Fig. 8B). Histograms of the population distribution of ODI values between control mice and those receiving 14 of early postnatal hypoxia were also nearly identical (Fig. 8C).

Binocular neurons have orientation tuning for each eye that refines during the critical period of visual development (P21-P32) to become more closely matched for preferred orientation. At P20, the average difference in preferred orientation for binocular neurons is near 30 degrees, but by P32 and into adulthood the difference in binocular matching of orientation is less than 20 degrees (Brown & McGee 2023; Tan et al. 2020; Wang et al. 2010). The matching of preferred orientation was similar for normoxic mice and 14d hypoxia mice (Fig. 8 D,E). Thus, while SF tuning is perturbed by 14d hypoxia, several measurements of orientation tuning were normal (Figs. 7 and 8).

## Discussion

CVI is a severe and increasingly prevalent developmental visual disorder (Hatton et al. 2007; Williams et al. 2021). The lack of an animal model is a barrier to progress for both understanding the circuit basis of visual deficits associated with CVI and developing effective treatments. Clinical studies have revealed that CVI is closely linked to perinatal hypoxia, pre-term birth, cerebral palsy, and PVL (Khetpal & Donahue 2007). Here we assessed whether established rodent hypoxia models for PVL could be adapted to yield a mouse model of CVI (Ment et al. 1998; Turner et al. 2003; Weiss et al. 2004).

Testing increasing durations of early postnatal hypoxia revealed that 14d was sufficient to produce serious deficits in binocular depth perception and visual acuity without compromising gross motor function or retinal function (Figs. 2 and 3). The visual impairments associated with CVI are not confined to stereopsis and acuity, but deficits in either are frequent (Sakki et al. 2021). In fact, acuity is a principal component for stratifying or subgrouping patients with CVI (Jimenez-Gomez et al. 2022; Sakki et al. 2021). Mice receiving 14d of early postnatal hypoxia also displayed a range of deficits in acuity and depth perception ranging from normal acuity (.4-.5 cpd) to severe impairments (<.18 cpd). While no animal model can be expected to capture every facet of a complex human visual disorder, particularly one with a diverse etiology, we conclude that 14d of early postnatal hypoxia provides an effective mouse model of CVI related to perinatal hypoxia and PVL.

Comparing cohorts of normoxic mice and 14d hypoxia mice yielded intriguing findings. Overall, the organization of retino-fugal projections was normal as innervation patterns to the SCN, OPN, SC, and dLGN were unremarkable and similar to normoxic controls (Fig. 4). However, the dLGN was smaller in mice receiving 14d hypoxia and the percent overlap of retinogeniculate projections more variable (Fig. 5). Interestingly, a recent clinical imaging study has demonstrated that patients with CVI associated with PVL have significantly smaller thalamic volumes than either preterm controls or patients with CVI not associated with PVL (Drottar et al. 2025).

In V1, the orientation tuning, binocular tuning, and binocular matching of orientation preference were normal overall for excitatory neurons in L2/3. In contrast, the distribution of preferred SFs tuning was shifted to slightly lower values relative to normoxic mice, and there were fewer neurons responsive to the higher SFs near normal acuity thresholds. This compression of SF tuning may be the underlying mechanism for lower acuity displayed by many 14d hypoxia mice.

Future work may assess whether this mouse model also disrupts other facets of visual function including affected in CVI including visual attention (Wang & Krauzlis 2018), oculomotor dysfunction (Michaiel et al. 2020; Tabata et al. 2011), and/or reduced contrast sensitivity (Histed et al. 2012). Multiple measurements of visual function and neuronal function performed in the same mice would better triangulate visual impairments with perturbations to central visual circuitry. We propose this mouse model will provide a framework for exploring therapeutic interventions to treat CVI.

## Methods

**Table 1.** Reagents and resources employed in the study.

| REAGENT or RESOURCE | SOURCE | IDENTIFIER |
| --- | --- | --- |
| **Deposited data** |  |  |
| Measurements from behavioral assays |  | N/A |
| Calculations of thalamic volume and tracing |  | N/A |
| Calculated tuning properties for all neurons |  | N/A |
| **Experimental models: Organisms/strains** |  |  |
| Mouse: B6;DBA-Tg(tedO-GCaMP6s)2Niell/j | The Jackson Laboratory | RRID: ISMR_JAX: 024742 |
| Mouse: B6;Cg-Tg(Camk2a-tTA)1Mmay/DboJ | The Jackson Laboratory | RRID: ISMR_JAX: 007004 |
| Mouse: Crl:CD1(ICR) | Charles River | RRID:IMSR_CRL:022 |
| **Software and algorithms** |  |  |
| MATLAB | Mathworks | https://www.mathworks.com/ |
| Processing2 | Processing | https://processing.org/ |
| **Reagents** |  |  |
| Cholera Toxin Subunit B 488 conjugate, Cat# C22841 | Invitrogen |  |
| Cholera Toxin Subunit B 555 conjugate, Cat# C34776 | Invitrogen |  |

### Lead contact

Further information and requests for resources and reagents should be directed to and will be fulfilled by the Lead Contact, Aaron McGee.

### Data availability

The Deposited data listed in the table above have been deposited in Mendeley data at:

### Experimental model and subjects

All procedures were approved by University of Louisville Institutional Animal Care and Use Committee and were in accord with guidelines set by the US National Institutes of Health. Mice were anesthetized by isoflurane inhalation and euthanized by carbon dioxide asphyxiation or cervical dislocation following deep anesthesia in accordance with approved protocols. Mice were housed in groups of 5 or fewer per cage in a 12/12 light dark cycle. Animals were naive subjects with no prior history of participation in research studies.

### Mice

Pups were fostered to CD-1 dams (Charles River). Calcium imaging was performed on mice expressing GCaMP6S in excitatory neurons in forebrain. The *CaMKII-tTA* (stock no. 007004) and *TRE-GCaMP6s* (stock no. 024742) transgenic mouse lines were obtained from Jackson Labs (Mayford et al. 1996; Wekselblatt et al. 2016). All other experiments were performed on mice from crosses of *CaMKII-tTA* and *TRE-GCaMP6s* that lacked one of the two transgenes. Mice were genotyped with primer sets suggested by Jackson labs.

### Hypoxia exposure

C57Bl/6J pups were fostered to CD-1 dams at P2 (Turner et al. 2003). At P3, CD-1 dams with their C57BL6 pups were transferred to custom-designed chambers (Oxycycler model A84XOV, BioSpherix, Lacona, NY) chambers in which the ambient O_2_ levels were maintained at 9.5% ± 1% by automated addition of nitrogen gas (Ment et al. 1998; Turner et al. 2003; Weiss et al. 2004). Following 7 days (P11), 14 days (P17), or 30 days (P33) days, of hypoxia, the mice were returned to standard housing and weaned at approximately P28.

### Rotarod task

Testing on the rotarod was adapted from our published method as follows (Park et al. 2014). Adult mice (>P45) were handled for 3-5 minutes each. On the same day, mice were placed on the stationary rod on the rotarod (IITC Life Science, Woodland Hills, CA) for 5 minutes to acclimate to the equipment. The following day, mice spent 5 more minutes acclimating to the rod. For the following 3 days, mice were tested on the rotating rod and 5 trials were performed a day as the rod increased in speed (4 to 44 rpm over 2 minutes) until the mouse fell from the rod to a platform below. Motor performance was recorded as time spent on accelerating rod for each trial (Jones & Roberts 1968).

### Pole descent visual cliff task (PDCT)

Testing on the PDCT was in accordance with our published method (Boone et al. 2021). In brief, adult mice had their whiskers shaved and were acclimated to handling for 3-5 minutes for each of 3 days before testing. During testing, mice were placed at the top of a pole perpendicular to a small cone sitting on thin metals rails above a glass surface. Below the glass, the interior of the testing chamber is covered in a black and white checkerboard pattern (2.5cm square). Three quadrants of a checkerboard surface parallel to the glass were positioned 15.2 cm below the glass surface and one quadrant was positioned 2.5cm below the glass. Mice were tested in interleaved trials and the position of the quadrant closest to the glass positioned at random for each trial by rotating the testing chamber. Mice were tested in 10-15 trials. Performance was measured as the fraction of trials that mice exited the pole above the quadrant that was nearest to the glass surface.

### Visual water task

Testing on the visual water task was in accordance with our method based on published work (Brown et al. 2024; Prusky et al. 2000; Stephany et al. 2014). In brief, mice were first trained to swim towards a platform associated with a static grating of 0.1 cycles per degree (cpd) until they could perform the task with at least 80% accuracy for 20 consecutive trials. Following this binocular training, mice received a monocular lid suture for the left eye and re-trained until the task was completed with at least 80% accuracy for 20 consecutive trials with monocular vision. Following monocular training, mice began monocular acuity testing at 0.1 cpd. The spatial frequency of the stimulus was increased by ∼0.025 cpd with the successful completion of 3 consecutive trials. After one failure, mice would then be required to complete the task accurately for 5 additional trials at the same spatial frequency and subsequent spatial frequencies tested. A second failure at the same spatial frequency requires mice to complete a total of 10 trials from the start of testing at that spatial frequency with no additional mistakes to progress to the next spatial frequency. Spatial frequency at failure (3 misses) was recorded for 3 or more rounds of testing (Stephany et al. 2014). Visual acuity is reported as the average of the 3+ spatial frequencies at which mice failed to reach passing criteria. The monocular lid suture was removed following testing.

### Anterograde labeling of retinofugal projections

Eye injections with Cholera toxin B (CTB) conjugated to Alexa Fluor dyes was performed as described in our published methods (Demas et al. 2006; Dilger et al. 2015; Jaubert-Miazza et al. 2005). In brief, mice were anesthetized by isoflurane (4% induction, 1% to 1.5% maintenance) following subcutaneous carprofen (5 mg/kg). The sclera was punctured using a sharp-tipped glass pipette, and some vitreous was removed. Another pipette, containing a 0.1–0.2% solution of the B subunit of cholera toxin (CTB, ThermoFisher Scientific) conjugated to either Alexa Fluor 555 (orange) or Alexa Fluor 488 (green), dissolved in distilled water, was introduced into the opening created by the initial pipette. The 5-8 μl CTB was injected using a Picospritzer (Parker-Hannifin). The following day, mice received a second dose of carprofen. Each eye received an injection of a different fluorescent conjugate to enable simultaneous and independent visualization of retinal terminal fields originating from each eye.

### Histology

Brain tissue was harvested 3–4-days after anterograde labeling of retinofugal projections. To collect brain tissue, mice were deeply anesthetized with isoflurane and transcardially perfused with phosphate buffered saline (PBS) followed by 4% paraformaldehyde in PBS. The brains were removed and post-fixed overnight at 4° C. Brains were sectioned in the coronal plane at 70 μm using a vibratome, then mounted serially, and coverslipped with Prolong Gold (Invitrogen). Epifluorescence microscopy was performed with a BX-51 upright microscope (Olympus) and Retiga EX B 12-bit monochrome camera. Images were taken through a 10X 0.25NA PLAN objective. Representative fluorescent images of the suprachiasmatic nucleus (SCN), the dorsolateral geniculate nucleus (dLGN), the olivary pretectal nucleus (OPN), and the superior colliculus (SC) were acquired and digitized separately using a BrightLine TxRed-A-Basic-OMF filter cube (Semrock) to image Alexa Fluor 555 and a BrightLine FTIC-A-Basic-OMF filter cube (Semrock) to image Alexa Fluor 488.

### Image analysis of eye specific projections in dLGN

We quantified the spatial extent of retinogeniculate projections arising from each eye by measuring the amount of area their domains occupied in dLGN (Demas et al. 2006; Dilger et al. 2015; Jaubert-Miazza et al. 2005). To determine the spatial extent of labeled retinal projections in dLGN, gray-scale images were converted into binary images using a threshold of 10 percent. Using the binarized images, we measured the spatial extent of ipsilateral and contralateral projections by counting the total number of labeled pixels within the boundaries of the dLGN. To determine the extent to which projections from the two eyes overlap, we superimposed the binarized images and counted the pixels that contained both signals. The size of ipsilateral and contralateral projections were measured, averaged across three successive sections through the middle of the dLGN, and expressed as a percentage of the total area of the dLGN. The size of overlapping terminal fields was reported as a percentage of the size of the ipsilateral projections.

### Full-Field Electroretinogram (ffERG)

Mice were dark-adapted overnight, anesthetized with a solution of ketamine (80mg/kg) and xylazine (16 mg/kg) administered via intraperitoneal injection (IP), and prepared for ffERG recordings under dim red light (Brown et al. 2024). Pupils were dilated and accommodation relaxed with the application of eye drops, with 2.5% phenylephrine hydrochloride Ophthalmic Solution (Bausch+Lomb, NDC 82260-102-10) and 0.5% Tropicamide Ophthalmic Solution (Bausch+Lomb NDC 24208-590-64) for 30 seconds. Eyes were then rinsed three times with sterile irrigating solution (Alcon Balanced Salts Solution [BSS]). A contact lens with a gold electrode (LKC Technologies Inc.) was placed on the cornea and position maintained with a hypromellose ophthalmic solution (Vista Gonio Eye Lubricant). Ground and reference needle electrodes were placed in the tail and on the midline of the forehead, respectively. Body temperature was maintained using a feedback-controlled electric heating pad (37°C).

Scotopic responses were measured at two test flash intensities (0.025 and 0.05 cd s/m^2^). After 10 minutes of adaptation to a rod-saturating background (20 cd/m^2^). Photopic responses were measured at two test flash intensities (10 and 100 cd s/m^2^). For each stimulus, a-wave, b-wave amplitudes, and b-wave implicit times were analyzed with a custom MATLAB code. The a-wave was measured from baseline (recorded 10-20 msec before stimulation) to the negative trough. The b-wave was measured from the a-wave trough to the b-wave peak. The flash intensities in the protocol for this experiment, scotopic flash (0.025 cd s/m^2^) and the photopic flash (100 cd s/m^2^), matched the visual water task screen luminance, allowing for direct comparison between behavioral vision acuity and retinal function. ERGs were performed within 2 weeks after concluding measuring acuity with the visual water task.

### Cranial windows

Wide field epi-fluorescent calcium imaging and two-photon calcium imaging were performed though a cranial window as previously described (Brown & McGee 2023; Trachtenberg et al. 2002). In brief, mice were administered carprofen (5 mg/kg) and buprenorphine (0.1 mg/kg) for analgesia and anesthetized with isoflurane (4% induction, 1% to 2% maintenance). The scalp was shaved and mice were mounted on a stereotaxic frame with palate bar and their body temperature maintained at 37°C with a heat pad controlled by feedback from a rectal thermometer (TCAT-2LV, Physitemp). The scalp was resected, the connective tissue removed from the skull, and a custom aluminum headbar affixed with C&B metabond (Parkell). A circular region of bone 3 mm in diameter centered over left visual cortex was removed using a high-speed drill (Foredom). Care was taken to not perturb the dura. A sterile 3 mm circular glass coverslip was sealed to the surrounding skull with cyanoacrylate (Pacer Technology) and dental acrylic (Ortho-jet, Lang Dental). The remaining exposed skull likewise sealed with cyanoacrylate and dental acrylic. Mice recovered on a heating pad and returned to standard housing for at least 2 days prior to 2-photon imaging.

### Wide field epi-fluorescent calcium imaging

After implantation of the cranial window and before 2-photon imaging, the binocular zone of visual cortex was identified with wide field calcium imaging similar to our method for optical imaging of intrinsic signals adapted from published studies (Frantz et al. 2016; Kalatsky et al. 2005). In brief, mice were anesthetized with isoflurane (4% induction), provided a low dose of the sedative chlorprothixene (0.5 mg/kg IP; C1761, Sigma) and secured by the aluminum headbar. The eyes were lubricated with a thin layer of ophthalmic ointment (Puralube, Dechra Pharmaceuticals). Body temperature was maintained at 37°C with heating pad regulated by a rectal thermometer (TCAT-2LV, Physitemp). Visual stimulus was provided through custom-written software (MATLAB, Mathworks). A monitor was placed 25 cm directly in front of the animal and subtended +40 to −40 degrees of visual space in the vertical axis. A horizonal white bar (2 degrees high and 20 degrees wide) centered on the zero-degree azimuth drifted from the top to bottom of the monitor with a period of 8 seconds. The stimulus was repeated 60 times. Cortex was illuminated with blue light (475 ± 30 nm) (475/35, Semrock) from a stable light source (Intralux dc-1100, Volpi). Fluorescence was captured utilizing a green filter (HQ620/20) attached to a tandem lens (50 mm lens, Computar) and camera (Manta G-1236B, Allied Vision). The imaging plane was defocused to approximately 200 μm below the pia. Images were captured at 10 Hz as images of 1,024 × 1,024 pixels and 12-bit depth. Images were binned spatially 4 × 4 before the magnitude of the response at the stimulus frequency (0.125 Hz) was measured by Fourier analysis.

### Visual stimuli and two-photon calcium imaging

Visual stimulus presentation and image acquisition were both performed according to our published methods which were modified from published studies (Brown et al. 2024; Brown & McGee 2023; Jimenez et al. 2018; Tan et al. 2020). In brief, a battery of static sinusoidal gratings was generated in real time with custom software (Processing, MATLAB). Stimulus presentation was synchronized to the imaging data by time stamping the presentation of each visual stimulus to the image acquisition frame number a transistor– transistor logic (TTL) pulse generated with an Arduino at each stimulus transition. Orientation was sampled at equal intervals of 15 degrees from 0 to 150 degrees (12 orientations). SF was sampled in 10 steps on a logarithmic scale at half-octaves from 0.028 to 1.02 cpd. An iso-luminant grey screen was included (blank) was provided as a ninth step in the SF sampling as a control. Spatial phase was equally sampled at 45-degree intervals from 0 to 315 degrees for each combination of orientation and SF. Gratings with random combinations of orientation, SF, and spatial phase were presented at a rate of 4 Hz on a monitor with a refresh rate of 60Hz. Imaging sessions were 15-20 minutes (3600-4800 gratings presented in total). Consequently, each combination of orientation and SF was presented 33 times on average (range 17 to 56) for a 20-minute session. The monitor was centered on the zero azimuth and elevation 35 cm away from the mouse and subtended 45 (vertical) by 80 degrees (horizontal) of visual space.

Imaging was performed with a resonant scanning 2-photon microscope controlled by Scanbox image acquisition and analysis software (Neurolabware). The objective lens was fixed at vertical for all experiments. Fluorescence excitation was provided by a tunable wavelength infrared laser (Ultra II, Coherent) at 920 nm. Images were collected through a 16× water-immersion objected (Nikon, 0.8 NA). Images (512 × 796 pixels, 520 × 740 μm) were captured at 15.5 Hz at depths between 150 and 400 μm. Eye movements and changes in pupil size were recorded using a Dalsa Genie M1280 camera (Teledyne Dalsa) fitted with 50 mm 1.8 lens (Computar) and 800 nm long-pass filter (Edmunds Optics). Imaging was performed on alert mice positioned on a spherical treadmill by the aluminum head bar affixed to the skull. The visual stimulus was presented to the contralateral eye by covering the fellow eye with a small custom occluder.

### Image Processing

Image processing was performed as described previously (Brown & McGee 2023; Tan et al. 2020). Imaging series for each eye were motion corrected with the SbxAlign tool. Regions of interest (ROIs) corresponding to excitatory neurons were selected manually with the SbxSegment tool following computation of pixel-wise correlation of fluorescence changes over time from 350 evenly spaced frames (∼1%). ROIs for each experiment were determined by correlated pixels the size similar to that of a neuronal soma. The fluorescence signal for each ROI and the surrounding neuropil were extracted from this segmentation map.

### Image Analysis

Image analysis was performed as described previously with minor modifications (Brown & McGee 2023). The fluorescence signal for each neuron was extracted by computing the mean of the calcium fluorescence within each ROI and subtracting the median fluorescence from the surrounding perimeter of neuropil (Ringach et al. 2016; Tan et al. 2020). An inferred spike rate (ISR) was estimated from adjusted fluorescence signal with the Vanilla algorithm (Berens et al. 2018). A reverse correlation of the ISR to stimulus onset was used to calculate the preferred stimuli (Brown & McGee 2023; Jimenez et al. 2018; Ringach et al. 2016; Tan et al. 2020). Neurons that satisfied 3 criteria were categorized as visually responsive: (1) the ISR was highest with the optimal delay of 4 to 9 frames following stimulus onset. This delay was determined empirically for this transgenic GCaMP6s mouse (Brown & McGee 2023; Tan et al. 2020); (2) the SNR was greater than 1.3. The signal is the mean of the spiking standard deviation at the optical delay between 4 and 9 frames after stimulus onset and the noise this value at frames −2 to 0 before the stimulus onset or 15 to 18 after it (Jimenez et al. 2018; Tan et al. 2020). (3) and neuron responded to at least 13% of the presentations of the preferred stimulus. Visual responsiveness for every neuron was determined independently for each eye. The visual stimulus capturing the preferred orientation and SF was the determined from the matrix of all orientations and SFs presented as the combination with highest average ISR.

The preferred orientation for each neuron was calculated as:

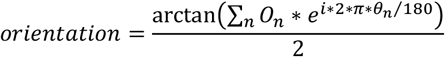

O_n_ is a 1 × 12 array of the mean z-scores associated with the calculation of the ISR at orientations Q_n_ (0 to 150 degrees, spaced every 15 degrees). Orientation calculated with this formula is in radians and was converted to degrees. The tuning width was the full width at half-maximum of the preferred orientation.

The preferred SF for each neuron was calculated as:

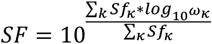

Sf_k_ is a 1 × 10 array of the mean z-scores at SFs w_k_ (10 equal steps on a logarithmic scale from 0.028 to 1.02. cpd). Tails of the distribution were clipped at 25% of the peak response. The tuning width was the full width at half-maximum of the preferred SF in octaves. The percent visually responsive neurons with significant responses at each SF was determined by comparing the distribution of ISR values at each SF versus the stimulus blank with a KW-test with Dunn’s correction for 10 comparisons. Neurons with P < 0.01 for a given SF were considered significant responses at that SF (Salinas et al. 2017).

### Statistics

Sample size was not predetermined based on statistical methods. Statistical analyses were conducted utilizing Prism 8 software (GraphPad Software). All data sets were tested for normality. Parametric tests, such as the Brown Forsythe and Welch ANOVA test and Welch’s t-test, were employed for data conforming to a normal distribution. Nonparametric tests including the Mann–Whitney (MW) test for pairwise comparisons, the Kruskal–Wallis (KW) test with Dunn’s correction for multiple comparisons were utilized for data that did not conform to a normal distribution. The repeated measure two-way mixed model with Geisser-Greenhouse correction was used for the rotarod results instead of an analysis of variance due to missing data points.

## Acknowledgements

This work was supported by a grant from the National Eye Institute (R01EY034092 to AWM) and a Jewish Heritage Fund for Excellence Research Enhancement Grant (WG). Schematics for the hypoxia exposure paradigm, the rotarod, and the PDCT, were designed and/or modified from Biorender under the license held by the Department of Anatomical Sciences and Neurobiology at the University of Louisville.

## Author contributions

DKO, CAA, JC, WG, and AWM conceived and designed the study. DKO, CAA, and JC, performed experiments. DKO, CAA, JC, WG, and AWM analyzed the data. DKO, WG, and AWM wrote the manuscript.

## Competing interests

The authors have declared that no competing interests exist.

## Notes

### Competing Interest Statement

The authors have declared no competing interest.

